# Ribosomal biogenesis factor links ribosome homeostasis to host relevant stress survival in *Aspergillus fumigatus*

**DOI:** 10.64898/2026.10.25.761613

**Authors:** John Adeyemi Adeoye, Hilla Hayby, Neta Shlezinger

## Abstract

The inhalation of *Aspergillus fumigatus* spores into the human pulmonary tract places conidia in direct contact with host immune cells which deploy a coordinated killing mechanism aimed at conidia clearance and preventing invasive infection. The fungus also employs cellular processes that aid its survival during these host mediated stresses, which allows the conidia to sustain growth while resisting attacks by the host immune cells. Ribosomal biogenesis plays a vital role in growth, survival, and stress adaptation in eukaryotic cells. In filamentous fungal pathogens, this process supports the transition of inhaled spores into invasive hyphae and is therefore an important aspect of fungal pathogenicity. It is, however, unclear whether ribosomal biogenesis factors contribute to stress survival beyond their canonical ribosomal roles and provision of support for vegetative growth. In this study, we investigated the role of *A. fumigatus* ribosomal biogenesis factor BFR2 in contribution to fungal stress survival and resistance to neutrophil killing. In agreement with its ribosomal biogenesis role, the repression of BFR2 impairs 18S ribosomal rRNA formation, and this impairment directly extends to the inhibition of germination and vegetative growth of fungal colonies. We optimized an “induction/shut-off” system that allows the transient expression of BFR2 in swollen conidia followed by subsequent depletion of BFR2 to investigate its role in stress response outside of its ribosomal biogenesis role. This transient induction followed by shut-off did not result in detectable reduction in ribosomal RNA abundance, global translation capacity, and biomass accumulation therefore indicating that pre-induction of BFR2 supports normal growth physiology in *A. fumigatus* over the duration tested. Nevertheless, the depletion of BFR2 following a transient induction increased *A. fumigatus* susceptibility to oxidative, replication and DNA-damaging stress while preserving tolerance to endoplasmic reticulum (ER) stress. The depletion of BFR2 under these conditions also increased *in vitro* neutrophil mediated killing without altering conidia uptake. In a mouse model of invasive aspergillosis, BFR2 depletion reduced conidial survival within neutrophils and pulmonary fungal burden, and in an immunocompromised mice model, the depletion of BFR2 attenuated virulence thereby extending survival by a few days. These findings identify BFR2 as a crucial link between ribosomal biogenesis and fungal stress survival during host relevant stresses. Thus, targeting ribosomal biogenesis factors that underpin stress survival represents a promising strategy for antifungal development that could target both ribosomal biogenesis function and also stress response functions.

## Introduction

Invasive aspergillosis, caused predominantly by airborne fungal pathogen, *Aspergillus fumigatus*, represents one of the most lethal fungal infections largely affecting immunocompromised individuals, yet effective therapeutic options are critically constrained to three classes of antifungals^1–5^. With over 2 million annual cases of infection globally, and a mortality rate ranging from 40% - 90% in high-risk groups including patients with hematological malignancies, patients receiving prolonged corticosteroid therapy, and solid organ transplant recipients even when first-line antifungal therapy is initiated promptly^6^. The therapeutic effectiveness of current classes of antifungal drugs against *A. fumigatus* is further challenged by the emergence of drug-resistant *A. fumigatus* isolates which have been isolated with increasing frequency across multiple continents and are partially associated with cases of treatment failure in the clinic^7–11^. The mechanistic basis of current antifungal agents converges on a set of fungal cell wall and cell membrane integrity targets. Echinochandins inhibit β-(1,3)-glucan synthase, azoles and polyenes disrupt ergosterol biosynthesis, and polyenes such as Amphotericin B directly binds ergosterol in fungal membranes to disrupt membrane integrity^12–18^. This convergence leaves other attractive therapeutic targets such as the fungal stress response network, metabolic regulatory networks, and cell survival pathways largely unexploited. As resistance mechanisms to current antifungal arsenal accumulate alongside an expanding population of immunocompromised patients population, the urgency to identify novel, essential fungal proteins with roles in stress adaptation and survival increases becomes both desirable and necessary for the future development of therapeutic options. Expanding the landscape of druggable targets therefore requires a systematic characterization of proteins that integrate stress adaptation and ability to survive within hostile host environment with core cellular function.

Upon the inhalation the stress response pathways of *A. fumigatus* conidia are tested during this early window of infection at the host-pathogen interface such that they are subject to a coordinated and chemically hostile host immune environment. Phagocytes such as alveolar macrophages and recruited neutrophils are the first-line immune cell defense against *A. fumigatus* conidia^2,19,20^. These phagocytes deploy NADPH oxidase-generated reactive oxygen species (ROS) within the phagosomal compartment producing an oxidative microenvironment capable of inflicting damage across macromolecular targets such as lipid peroxidation, protein carbonylation, and direct DNA strand breaks, thus serving as their primary cytotoxic mechanism of achieving killing^21–25^. Neutrophil-derived ROS constitute the primary genotoxic threat, making oxidative stress tolerance a genuine determinant of pathogen survival. A recent study demonstrated that conidia internalized by neutrophils exhibit DNA fragmentation patterns consistent with regulated cell death (RCD)-like processes in a manner that is consistent with ROS generation within the phagolysosome^26^. *A. fumigatus* conidia are also able to withstand this ROS-induced genotoxic progression implying the presence of an active stress response network capable of modulating these RCD-like outcomes^26^. Understanding how the fungus senses, tolerates and resists oxidative stress-induced killing by neutrophils is therefore relevant and important for exploiting vulnerabilities in *A. fumigatus* stress response and survival machinery that could be pharmacologically exploited.

Ribosomal biogenesis has recently emerged as a dynamic hub that integrates stress tolerance and coordinates cell survival decisions in eukaryotic cells^27–33^. Hence, it is positioned as an attractive, underexplored and non-classical therapeutic target axis in fungal pathogens. In mammalian systems, the ribosomal biogenesis factors do not only play a role as constitutive housekeeping proteins, but they also participate in dynamic regulatory processes that reprogram the cell in response to genotoxic stressors to arbitrate cell-fate decisions between survival and death^30,34,35^. Ribosomal biogenesis stress in mammalian cells is usually triggered by DNA damage, nutrient deprivation or oxidative assault^36,37^. These stresses activate a non-transcriptional nucleolar stress surveillance pathway in which specific ribosomal biogenesis factors are released from the nucleolus, bind to MDM2, and stabilize p53 to coordinate cell cycle arrest and apoptosis^38–40^. The protein AATF (Apoptosis Antagonizing Transcription Factor), originally identified as a component of the RNA polymerase II transcription machinery and subsequently found to be essential for 18S rRNA processing and small subunit (SSU) assembly, occupies a critical node at the intersection of ribosome biogenesis, DNA damage response and cell survival^41–47^. AATF depletion in mammalian cells impairs SSU maturation, sensitizes cells to apoptosis induced by genotoxic stressors, and the complete knock-out of AATF in mice results in non-viable embryo at the blastocyst stage, thereby establishing its essentiality for cellular viability under replicative stress^46,48,49^. In *Saccharomyces cerevisiae*, BFR2, the functional homolog of AATF, is essential for viability and required for pre-18s rRNA processing and SSU assembly. Its depletion produces a lethal effect in small ribosomal subunit biogenesis phenotypically mirroring AATF loss in mammalian embryo^50,51^. The conservation of AATF function across metazoans and budding yeasts at both structural and stress-regulatory level indicates an evolutionary ancient, functionally constrained node that couples the cell’s translational capacity to its stress tolerance. Rather than passively supporting translation, BFR2 appears to possibly play a role where biosynthetic capabilities and stress tolerance are jointly controlled. This is particularly important in *A. fumigatus* because by retaining the stress response function observed in AATF, BFR2 would simultaneously control two processes that are critical during infection - the biosynthetic capacity required for vegetative growth, and stress tolerance required to survive host-relevant stresses.

In *A. fumigatus*, the transition from dormant, intrinsically stress resistant conidia to vegetatively active hyphae represent an important window in which both biosynthetic commitment and stress tolerance are simultaneously required. Thus, factors that govern this transition become attractive to therapeutic targets. Germination requires active reprogramming if the translational apparatus, rapid ribosomal accumulation, and simultaneous capacity to withstand stresses imposed by the environment^52,53^. Targeting a factor essential for ribosomal biogenesis during germination and that additionally modulates the fungal stress-responses to host-relevant stress would be doubly constraining on the fungi. On one hand, its loss will block growth even in absence of immune pressure, and on the other hand, its suppression would sensitize cells to immune killing. This dual function profile of BFR2 as the *A. fumigatus* functional homolog of stress-responsive ribosomal biogenesis factor AATF places it as a high-priority target. The role of BFR2 in *A. fumigatus* is mostly uncharacterized. Studies from the yeast model offer a glimpse into the ribosomal role of fungal BFR2 however, they are insufficient to resolve stress-regulatory roles in the context of filamentous fungal biology, infection relevant morphological transition, and host immune cell interactions^50^. Hence, stress biology of BFR2 is undefined at a mechanistic level in pathogenic filamentous fungi. Due to its essentiality in mammalian embryonic development and yeast viability, and its likely essentiality in *A. fumigatus*, its functional characterization requires a conditional expression system that allows the decoupling of its essential ribosome assembly role from any separable stress-regulatory function. A systematic characterization of BFR2 encompassing its essentiality, subcellular localization, ribosomal roles and function under a panel of host-relevant stresses is therefore required to determine whether BFR2 constitutes a dual function stress tolerance protein and hence, a viable therapeutic target.

To address this gap in knowledge, we deployed a conditional expression system using xylose-inducible promoter. We showed that by the repression of BFR2 in *A. fumigatus* arrests vegetative growth and germination and also produces a defect in 18S rRNA maturation consistent with SSU assembly failure. By bypassing BFR2 essentiality *via* transient induction, followed by depletion, we demonstrated that BFR2 is essential for conidia germination, vegetative growth and 18s rRNA processing, it also contributes to fungal stress survival, linking ribosomal biogenesis to adaptive responses during host infection. Together, these findings define BFR2 as a stress-responsive, essential ribosomal biogenesis factor with a previously unrecognized role in managing host-relevant stresses thereby positioning ribosomal biogenesis stress response networks as a mechanistically tractable and therapeutically relevant target in invasive aspergillosis.

## Results

### Nucleolar localized ribosomal biogenesis factor BFR2 is important for conidia germination, biomass accumulation and ribosomal biogenesis function in *Aspergillus fumigatus*

BFR2 is a 565-amino-acid protein in *A. fumigatus* that retains the AATF-Che1 leucine zipper-like domain and the AATF C-terminal (TRAUBE) domain found in its homologs in *H. sapiens* (560 aa), *A. oryzae* (519 aa), *A. lentulus* (566 aa), *C. albicans* (512 aa), *C. glabrata* (522 aa) and *S. cerevisiae* (534 aa) (Figure 1A). Pairwise sequence alignment showed that overall sequence identities between fungal BFR2 and mammalian AATF are low, at approximately 37%. Phylogenetic analysis placed *A. fumigatus* BFR2 within a fungal clade distinct from mammalian AATF clade (Figure S1A), identifying BFR2 as a distant fungal homolog of mammalian AATF.

**Figure 1.**
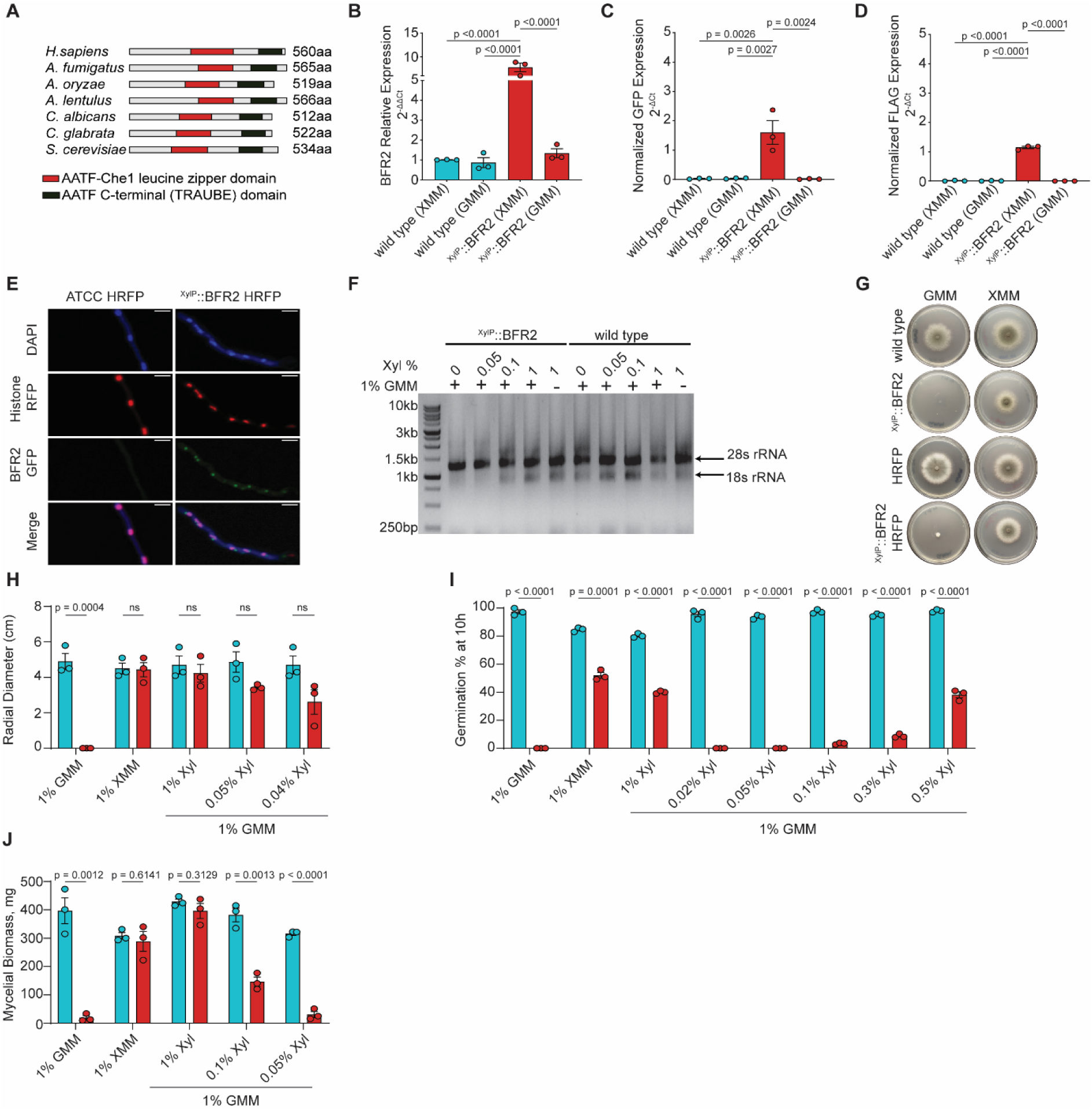
BFR2 is important for conidial germination, vegetative growth, biomass accumulation, and ribosome biogenesis function in *A. fumigatus*. (A) Domain organization of the AATF/Che-1 domain (red) and TRAUBE domain (black) in human AATF and BFR2 orthologs from *A. fumigatus*, *A. oryzae*, *A. lentulus*, *C. albicans*, *C. glabrata*, and *S. cerevisiae*. (B) BFR2 mRNA expression in wild type and ^XylP::^BFR2, following growth of 5 × 10⁶ conidia under fully inducing (1% XMM) or non-inducing (1% GMM) conditions at 37°C overnight. Expression was normalized to actin and is presented as mean relative BFR2 expression (2^-ΔΔCt^) ± SEM from three independent biological experiments. Statistical significance was assessed by one-way ANOVA with Tukey’s multiple comparisons test. Statistical significance was determined at p<0.05. (C–D) Relative expression of GFP (C) and FLAG (D), quantified as surrogate reporters to isolate construct-specific expression. Because GFP and FLAG transcripts are unique to the 3xFlag-BFR2-GFP cassette present in ^XylP::^BFR2 and absent from the wild-type genome, expression is reported as actin-normalized 2^-ΔCt^ values rather than 2^-ΔΔCt^, since wild type carries no GFP or FLAG sequence and therefore provides no baseline Ct for a ΔΔCt calculation. Data are presented as mean actin-normalized expression ± SEM from three independent biological experiments. Statistical significance was assessed by one-way ANOVA with Tukey’s multiple comparisons test. Statistical significance was determined at p<0.05. (E) Subcellular localization of BFR2. BFR2-GFP was expressed in the HRFP background strain, in which HRFP labels chromatin in the nucleus. HRFP and ^XylP::^BFR2-HRFP strains were grown at 37°C, and resulting hyphae were fixed and counterstained with DAPI to label nuclei. Imaging was performed by fluorescence microscopy; a representative image is shown (scale bar = 10μm). (F) Total RNA extracted from wild type and ^XylP::^BFR2 grown under a xylose titration series — fully repressing 1% GMM, 0.05% xylose in 1% GMM, 0.1% xylose in 1% GMM, 1% xylose in 1% GMM, and fully inducing 1% XMM — was resolved on a 1% agarose gel. 18S and 28S rRNA bands are shown. Gel image is representative of two independent experiments. (G–J) Essentiality of BFR2 for vegetative growth, germination, and biomass accumulation. (G) 1 × 10⁵ conidia from wild type, ^XylP::^BFR2, HRFP, and ^XylP::^BFR2-HRFP were inoculated at the centre of 1% GMM or 1% XMM agar plates and incubated at 37°C for 4 days. Representative images of vegetative growth are shown (three independent experiments). (H) Radial diameter of wild type and ^XylP::^BFR2 (1 × 10⁵ conidia) inoculated at the centre of agar plates containing fully repressing 1% GMM, fully inducing 1% XMM, or 1% GMM supplemented with 1%, 0.05%, or 0.04% xylose, incubated at 37°C for 4 days. Radial diameter was measured on day 4. (I) Germination of wild type and ^XylP::^BFR2 (5 × 10^5^ conidia) in fully repressing 1% GMM, fully inducing 1% XMM, or 1% GMM supplemented with 1%, 0.5%, 0.3%, 0.1%, 0.05%, or 0.02% xylose, incubated at 37°C for 10 hours. Germination was scored at 10 hours. (J) Biomass accumulation. 5 × 10⁷ conidia from wild type and ^XylP::^BFR2 were inoculated in 50 mL of fully repressing 1% GMM, fully inducing 1% XMM, or 1% GMM supplemented with 1%, 0.1%, or 0.05% xylose, and grown for 24 hours. Mycelium was harvested, lyophilized for 4 days, and weighed to determine mycelial biomass. For G–J: Data are presented as mean ± SEM from three biologically independent experiments. Statistical significance was assessed by Student’s t-test. Statistical significance was determined at p<0.05.

Three attempts using multiple independent targeting strategies to generate a complete BFR2 deletion mutant in *A. fumigatus* were unsuccessful, consistent with known BFR2 essentiality in yeast and for AATF in mammalian systems. A conditional expression strategy was therefore used in place of a full gene deletion, employing the xylose-inducible promoter system, which allows graded, dose-dependent control of gene expression based on xylose concentration in the growth medium and has been previously used to generate conditional mutants of essential genes in *A. fumigatus*. The *BFR2* disruption strain was generated by CRISPR-Cas9-mediated cleavage near the translational start codon, with non-homologous end joining introducing a premature stop codon which was confirmed by amplicon sequencing. This strain was complemented with a xylose-inducible 3xFlag-BFR2-GFP construct integrated ectopically at an unmapped genomic locus. Successful integration was confirmed by PCR of the full construct and detection of GFP expression; functional complementation was confirmed phenotypically. A 3xFlag epitope tag and GFP fluorescent marker were incorporated into the construct, enabling detection of BFR2 protein expression by western blotting and visualization of BFR2 localization by fluorescent microscopy.

BFR2 transcript abundance was quantified by RT-qPCR using primers targeting BFR2, normalized to actin and to wild type growing on 1% xylose minimal media as calibrator (Figure 1B). ^XylP::^BFR2 grown in XMM showed a 7.70-fold increase in BFR2 transcript relative to wild type grown in XMM and a 5.80-fold increase relative to ^XylP::^BFR2 grown in GMM. A consistent pattern was observed in the HRFP background (Figure S2A), where ^XylP::^BFR2-HRFP grown in XMM showed an 11.18-fold increase in BFR2 transcript relative to HRFP grown in XMM and a 3.37-fold increase relative to ^XylP::^BFR2-HRFP grown in GMM.

Since endogenous BFR2 transcript is still produced in the mutant background despite the premature stop codon, elevated signal in the BFR2-primer assay alone cannot be attributed unambiguously to the inducible construct. To isolate construct-specific expression, transcript levels of GFP and FLAG, both unique to the 3xFlag-BFR2-GFP cassette and absent from the wild-type genome, were quantified as surrogate reporters of xylose-driven BFR2 expression (Figures 1C–D). These are reported as actin-normalized 2^-ΔCt^ values rather than 2^-ΔΔCt^, since wild type carries no GFP or FLAG sequence and therefore provides no baseline Ct for a ΔΔCt calculation.

In the wild-type background, actin-normalized GFP levels reached 1.60 in ^XylP::^BFR2 under inducing conditions (XMM), dropping to 0.0134 in ^XylP::^BFR2 in non-inducing conditions (GMM), compared to 0.0274 and 0.0368 in wild type grown in inducing and non-inducing, respectively. FLAG levels showed the same pattern, reaching 1.15 in ^XylP::^BFR2 under inducing conditions and 0.00104 when grown in non-inducing conditions, compared to 0.00939 and 0.00713 in wild type in XMM and GMM, respectively.

The same pattern held in the HRFP background (Figure S2B–C). GFP levels reached 2.11 in ^XylP::^BFR2-HRFP under inducing conditions, and 0.00891 in non-inducing conditions, compared to 0.0197 and 0.0181 for HRFP in XMM and GMM, respectively. FLAG levels followed a similar pattern, reaching 1.33 in ^XylP::^BFR2-HRFP grown in inducing conditions and 0.000795 in non-inducing conditions, compared to 0.000190 and 0.000160 for HRFP in XMM and GMM, respectively.

Similarly, protein expression levels were substantially reduced, confirming that the conditional system achieves the intended repression and induction of BFR2 expression before any downstream phenotype was examined.

BFR2 fused to GFP expressed from the Xylose inducible construct in a Histone H2A::mRFP reporter background localized as discrete fluorescent foci at the periphery of each nucleus, rather than being distributed throughout the bulk chromatin (Figure 1E). Histone and Hoechst signal overlapped completely, as expected for the two markers of total DNA, whereas BFR2 fused to GFP was adjacent to, but did not overlap with HRFP or Hoechst-stained nuclear region.

Total RNA was extracted from wild type and ^XylP::^BFR2 mutants strains grown on 1% GMM, 1% XMM and 1% GM supplemented with 0.05%, 0.1% or 1% Xylose, and resolved by agarose gel electrophoresis. Wild type showed the expected 18S and 28S rRNA bands under every condition tested. (Figure 1D). In the ^XylP::^BFR2 mutants strain, the 18S rRNA band was completely absent on 1% GMM and on 1% GMM supplemented with 0.05% Xylose, and became detectable at 0.1% and 1% xylose, with band intensity relative to 28S rRNA increasing further at 1% xylose (Figure 1F).

Together, these observations show that BFR2 forms discrete foci adjacent to, but distinct from the bulk nuclear DNA in *A. fumigatus* which coincides with the subnuclear localization of nucleolus, and BFR2 expression is required for the formation of 18S rRNA.

Repression of BFR2 strongly impairs conidial germination. Our ^XylP::^ BFR2 conditional strain grown in complete repression condition (1% GMM, no xylose) produced essentially no germination at 10h, whereas graded addition of xylose produced a dose-dependent restoration of germination, with full induction yielding percentages about 50% of the wild type. These results indicate a critical threshold of BFR2 expression is required to initiate germination.

Given BFR2’s nucleolar localization and conservation of Che-1/TRAUB domains implicated in ribosomal biogenesis and RNA processing, it is possible explanation is that reduced BFR2 levels impair ribosomal biogenesis which in turn reduces overall protein synthesis and prevents the production of factors required to exit dormancy and initiate polarized growth. Lower concentrations of xylose that are unable to induce germination indicate that BFR2 expression level is below the threshold required for germination, intermediate xylose concentrations partially restore ribosomal biogenesis function of BFR2 resulting in partial germination and at full induction, ribosomal function is normalized to a near wild type function resulting in observed germination.

Repression of BFR2 expression caused severe vegetative growth defects on solid agar media. The ^XylP::^BFR2 mutant strain failed to produce any visible colony growth on 1% GMM after 4 days, whereas wild type grew robustly under the same condition (Figure 1E). Growth was fully rescued by 1% Xylose minimal media, where colony expansion of conditional mutants was comparable to wild type (Figure 1G). To determine the xylose threshold required to support growth, radial colony diameter was measured on 1% GMM supplemented with 1%, 0.05% or 0.04% xylose. Supplementation with 1% xylose restored growth to wild type levels, whereas supplementation with 0.05% or 0.04% xylose produced colonies that were visually reduced in diameter but were not significantly different from wild type by quantitative measurement (Figure 1H).

BFR2 expression was required for efficient conidial germination. Under complete repression conditions (1% GMM with no xylose), essentially no germination was detected at 10h post-inoculation, compared to 97.33% germination in the wild type (Figure 1I). Graded addition of xylose produced a dose-dependent restoration of germination: 8.82% at 0.3% xylose, 38.15% at 0. 5% xylose and 40% at 1% xylose, the latter reaching approximately 50% of wild type germination rate at 10h (Figure 1I), compared to 95.66% in wild type grown under the same conditions. Even at full induction, on 1% XMM, germination of ^XylP::^BFR2 mutant strain reached only approximately 52.03% of the wild type at 10h, compared to 84.66% in the wild type grown under the same conditions, indicating that xylose-driven expression does not fully reconstitute endogenous BFR2 levels.

Biomass accumulation in liquid culture mirrored the germination and vegetative growth phenotypes in a xylose concentration-dependent manner. The ^XylP::^BFR2 mutant strain grown in 1% GMM without xylose accumulated minimal biomass over 24h, whereas growth in 1% XMM produced biomass comparable to wild type over the same period (Figure 1H). Supplementation of 1% GMM with graded xylose concentrations produced a stepwise reduction in biomass accumulation: cultures supplemented wih 1% Xylose accumulated biomass comparable to wild type, whereas supplementation with 0.1% xylose reduced biomass accumulation by at least 66%, and supplementation with 0.05% xylose reduced biomass by approximately 90% relative to wild type at 24h (Figure 1J).

Taken together, these findings establish BFR2 as an essential regulator of growth and development in *A. fumigatus*. The nucleolar localization of BFR2 and its requirement for 18S rRNA biogenesis provide a molecular basis for the growth phenotypes observed upon its repression: failure to accumulate sufficient BFR2 protein abolishes 18S rRNA production, and this ribosomal biogenesis defect manifests as a coordinated failure across three independent growth readouts including conidial germination, vegetative colony expansion, and biomass accumulation in liquid culture, each of which responds to BFR2 induction in a graded, concentration-dependent manner. The agreement of these three phenotypes across identical xylose induction series argues that BFR2 expression level is the primary determinant of growth capacity, with a critical threshold below which the fungus cannot exit conidial dormancy or sustain vegetative growth. That even maximal xylose induction restores germination to only about 50% of wild type further indicates that the XylP promoter does not fully recapitulate endogenous BFR2 expression, and that the phenotypes described here represent a hypomorphic state whose severity would be exceeded by complete loss of BFR2 function.

### *A. fumigatus* sustains normal physiology upon transient BFR2 expression and subsequent depletion

To circumvent the germination defect observed under constitutive BFR2 repression, we established a two-phase experimental design in which BFR2 expression was first induced transiently for 6 hours in 1% XMM, which corresponds to the developmental window of conidial swelling in XMM, followed either by continued culture in 1% XMM or transfer to GMM to repress BFR2 expression(hereafter “shut-off condition”), with both conditions allowing progression into germlings over a further 6 hours (Figure 2A). This design permitted interrogation of the consequences of acute BFR2 depletion in cells that had already successfully initiated germination, independently of their earlier requirement for BFR2 during conidial activation.

**Figure 2.**
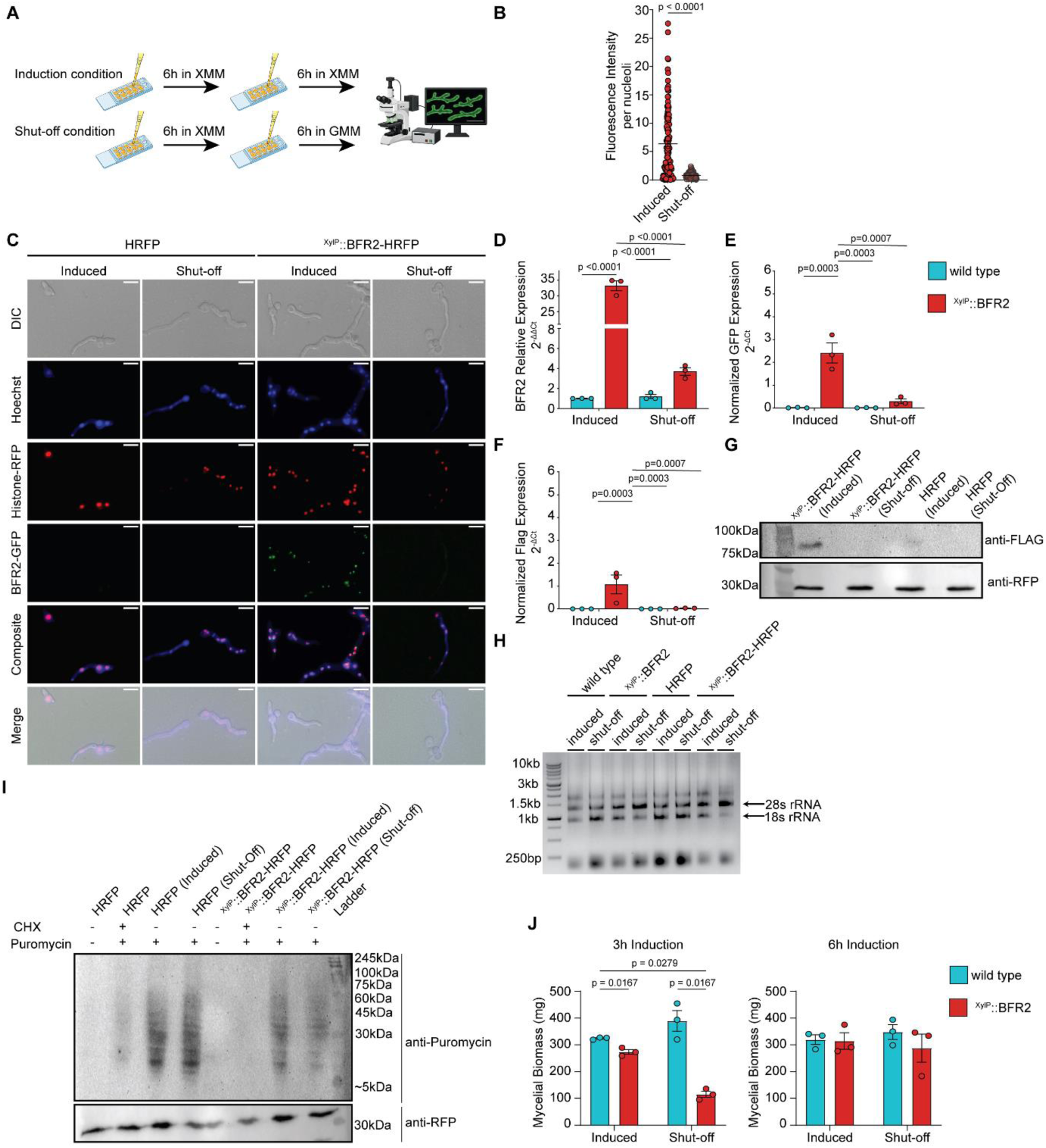
Transient expression followed by subsequent depletion of BFR2 sustains normal physiology in *A. fumigatus*. **(A)** Schematic of the induction/shut-off experimental design, developed to circumvent germination defects observed when BFR2 is not fully induced. BFR2 was pre-induced in 1% XMM (fully inducing conditions) for 6 hours, followed by a further 6-hour incubation in 1% GMM (shut-off) or continued 1% XMM (induced control) to initiate BFR2 depletion in the resulting germlings. For Panels B–H, cells were incubated for an additional 4 hours beyond this base protocol (16 hours of total culture time), matching the longest stress-exposure duration used in the assays presented in subsequent Figure 3, to confirm that BFR2 induction and depletion were sustained across the full-time window required for subsequent stress challenges. **(B–C)** Fluorescence intensity of BFR2-GFP over the post-induction window. **(B)** Quantification of BFR2-GFP expression across a 10-hour post-induction window (the 6-hour shut-off/continued-induction phase plus the additional 4-hour incubation described in A). **(C)** Fluorescence microscopy of HRFP and ^XylP::^BFR2-HRFP strains, imaged at the end of this window. Cells were labelled with Hoechst 33342 prior to imaging while HRFP labels nuclear chromatin. Panels are representative of two independent experiments (scale bar = 10 μm). **(D)** BFR2 mRNA expression in wild type and ^XylP::^BFR2 at the end of the protocol described in (A) (16 hours total), from 5 × 10⁶ conidia grown at 37°C. Expression was normalized to actin and is presented as mean relative BFR2 expression (2-ΔΔCt) ± SEM from three independent biological experiments. Statistical significance was assessed by one-way ANOVA with Tukey’s multiple comparisons test and determined at p<0.05. **(E–F)** Relative expression of GFP (E) and FLAG (F), quantified as surrogate reporters to isolate construct-specific expression, at the same endpoint as (D). Because GFP and FLAG transcripts are unique to the 3xFlag-BFR2-GFP cassette present in ^XylP::^BFR2 and absent from the wild-type genome, expression is reported as actin-normalized 2-ΔCt values rather than 2^-ΔΔCt^, since wild type carries no GFP or FLAG sequence and therefore provides no baseline Ct for a ΔΔCt calculation. Data are presented as mean actin-normalized expression ± SEM from three independent biological experiments. Statistical significance was assessed by one-way ANOVA with Tukey’s multiple comparisons test and determined at p<0.05. **(G)** Western blot analysis of BFR2 protein expression in the HRFP parental control and ^XylP::^BFR2-HRFP conditional mutant under induced and shut-off conditions. FLAG-tagged BFR2 was detected using an anti-FLAG antibody. RFP, detected using an anti-RFP antibody, served as the loading control. **(H)** Total RNA extracted from wild type, ^XylP::^BFR2, HRFP, and ^XylP::^BFR2-HRFP at the same endpoint as (D), resolved on a 1% agarose gel. 18S and 28S rRNA bands are shown. Gel image is representative of two independent experiments. **(I)** Translation efficiency of HRFP and ^XylP::^BFR2-HRFP at the same endpoint as (D). Conidia (5 × 10⁶) were grown under the respective conditions, then labelled with puromycin (15 μg/mL) for 30 minutes. A cycloheximide control (100 μg/mL) was included, added 30 minutes before the end of the incubation series and prior to puromycin addition; a no-puromycin control was also included. Germling cultures were then placed on ice to halt puromycin incorporation, harvested, and flash-frozen in liquid nitrogen. Protein was extracted, resolved by Western blot, and probed with anti-puromycin antibody (1:10,000). Blot shown is representative of two independent biological experiments. **(J)** Biomass accumulation under the induction/shut-off experimental set-up. Wild type and ^XylP::^BFR2 (5 × 10⁷ conidia) were incubated in their respective conditions (induced or shut-off) for 24 hours total: 6 hours of induction, followed by 6 hours of shut-off, then a further 12 hours in the same condition, allowing sufficient time for measurable mycelial growth to accumulate. As a comparison to justify selection of 6 hours as the standard induction time, a parallel condition was included in which BFR2 was induced for only 3 hours, before reaching the same 24-hour endpoint (left panel). Mycelium was harvested, flash-frozen, and lyophilized for 4 days, then weighed to determine mycelial biomass. Data are presented as mean ± SEM from three biologically independent experiments. Statistical significance was assessed by one-way ANOVA with Tukey’s multiple comparisons test and determined at p<0.05.

To validate the effectiveness of the two-phase experimental approach, the fluorescence intensity of ^XylP::^BFR2 fused to GFP was quantified by fluorescence microscopy coupled with quantitative image analysis over the 6h induction and the following 6h induction/shut-off window (Figure 2B, C). Cells maintained in 1% XMM sustained BFR2-GFP expression throughout the 12-h period, consistent with continued xylose-driven transcription during germling development. In contrast, cells transferred to 1% GMM exhibited a marked reduction in BFR2-GFP fluorescence intensity relative to the 1% XMM condition, consistent with downregulation of BFR2 following medium exchange.

RT-qPCR quantification of BFR2 transcript levels, reported as fold change (2^-ΔΔCt^) relative to the wild type or HRFP control, revealed the same pattern (Figure 2D). In the wild type background, BFR2 transcript levels increased 33.2-fold in the ^XylP::^BFR2 strain under inducing conditions (1% XMM), compared to a 1.01-fold change in the wild-type control. Following medium exchange to 1% GMM, BFR2 transcript levels in the ^XylP::^BFR2 strain declined to 3.71-fold, an 8.96-fold reduction from the induced state, while the wild-type control remained at 1.20-fold. The same pattern was observed in the HRFP background where BFR2 transcript levels increased 22.3-fold under inducing conditions and declined to 3.83-fold following shut-off, a 5.82-fold reduction, while the HRFP control remained at 1.00-fold (XMM) and 1.01-fold (GMM) (Figure S2A). In both genetic backgrounds, BFR2 transcript levels in the ^XylP::^BFR2 strain under shut-off conditions remained above the ∼1.0-fold background level of the respective control strain, rather than returning to it indicating some level of residual expression.

To confirm that these changes specifically reflected xylose-driven transcription of the 3xFlag-BFR2-GFP cassette rather than endogenous BFR2 transcript, GFP and FLAG transcript levels were also quantified in parallel as construct-specific surrogate reporters. As neither GFP nor FLAG sequence is present in the wild-type or HRFP genomes, these values are reported as actin-normalized 2^-ΔCt^ levels rather than fold change (Figure 2E-F). GFP transcript was near background in the wild-type and HRFP strains regardless of medium (2^-ΔCt^ = 0.0099 and 0.0038 in 1% XMM; 0.0050 and 0.0081 in 1% GMM, respectively) (Figure 2E-F, Figure S2B-C). In the ^XylP::^BFR2 strain, GFP transcript levels were 2.41 (wild-type background) and 3.56 (HRFP background) under inducing conditions, declining to 0.29 and 0.40 following shut-off, an 8.30-fold and 8.95-fold reduction, respectively (Figure 2E, Figure S2B). FLAG transcript levels followed the same pattern: near-background in the wild-type and HRFP strains (2^-ΔCt^ = 0.00035 and 0.00030 in 1% XMM; 0.00037 and 0.00032 in 1% GMM), compared to 1.07 (wild-type background) and 1.40 (HRFP background) in the induced ^XylP::^BFR2 strain, declining to 0.026 and 0.031 after shut-off, a 40.8-fold and 44.9-fold reduction (Figure 2F, Figure S2C). As with the BFR2 transcript data, GFP and FLAG signal in the shut-off condition did not return to the near-background levels of the wild-type and HRFP strains.

To assess whether acute BFR2 depletion following pre-induction compromises core cellular functions, 18S rRNA levels, translational capacity, and biomass accumulation was evaluated under shut-off conditions. Agarose gel electrophoresis of total RNA extracted after induction/shut-off revealed no detectable reduction in 18S rRNA levels of cells in shut-off condition relative to fully induced controls (Figure 2G). Translational capacity assessed by the SUnSET puromycin incorporation assay similarly revealed no significant impairment of global protein synthesis following 10 hours of BFR2 shut-off relative to induced controls (Figure 2H).

To determine the minimum pre-induction period required to support subsequent growth under shut-off conditions, mycelial biomass was quantified following either 3 or 6 hours of pre-induction in 1% XMM, each followed by transfer to 1% GMM for the remainder of a 20-hour experiment, with parallel cultures maintained continuously in XMM as controls. Following 3 hours of pre-induction, the ^XylP::^BFR2 mutant strain exhibited a pronounced reduction in biomass under shut-off conditions relative to both its induced counterpart and wild type controls (Figure 2I). Extending pre-induction to 6 hours fully restored biomass accumulation under shut-off conditions, with no significant differences detected between shut-off and induced ^XylP::^BFR2 mutant strain cultures or relative to wild type controls under either condition (Figure 2I). Wild type biomass accumulation was unaffected by medium exchange under either pre-induction regime (Figure 2I), confirming that the medium switch itself does not impose a growth penalty independent of BFR2 expression status.

Collectively, 18S rRNA abundance, global translational capacity, and biomass accumulation were each maintained following 6 hours of BFR2 pre-induction and subsequent depletion, establishing this two-phase regime as a valid experimental framework for interrogating BFR2 function independently of the early germination defect.

### Transient BFR2 expression followed by depletion renders *A. fumigatus* hypersensitive to oxidative, DNA damage, and replication stress

BFR2 depletion selectively impaired tolerance to replication stress and double-strand DNA damage in a manner dependent on the degree of BFR2 expression. To probe this, we challenged wild type and ^XylP::^BFR2 mutant strains with three distinct stressors on agar plates. Hydroxyurea (20mM), a ribonucleotide reductase inhibitor that depletes dNTP pools and stalls replication forks was used to induce replication stress, phleomycin (10µg/ml), a radio mimetic glycopeptide was used to induce double-strand DNA breaks, and DTT (20mM), a reducing agent that disrupts disulfide bond formation was used to elicit ER stress. On 1% XMM plates, which supports full BFR2 induction, the ^XylP::^BFR2 mutant strain grew comparably to wild type across all three conditions. Relative growth on hydroxyurea was 50.58% for ^XylP::^BFR2 versus 57.23% for wild type, on phleomycin it was 75.53% versus 93.43%, and on DTT it was 49.92% versus 50.77%; none of these differences reached statistical significance (Fig. 3A, top panel). By contrast, at the minimally inducing concentration of 0.05% xylose in 1% GMM, the lowest concentration of xylose identified to permit growth on agar plates, BFR2 depletion produced stressor-selective and severe growth defects. On hydroxyurea, relative growth of ^XylP::^BFR2 was significantly reduced to 14.58% compared to 49.20% for wild type (Figure 3A, bottom panel). On phleomycin, the reduction was even more pronounced, with ^XylP::^BFR2 achieving only 19.29% relative growth against 98.10% in wild type (Figure 3A, bottom panel), indicating near-complete sensitivity to double-strand DNA damage under conditions of partial BFR2 depletion. In contrast, DTT tolerance was fully preserved in ^XylP::^BFR2 at this xylose concentration (47.56% vs 56.26%), demonstrating that the growth defect is not a generalised consequence of stress exposure but reflects a specific requirement for BFR2 in the response to genotoxic and replication stress.

**Figure 3.**
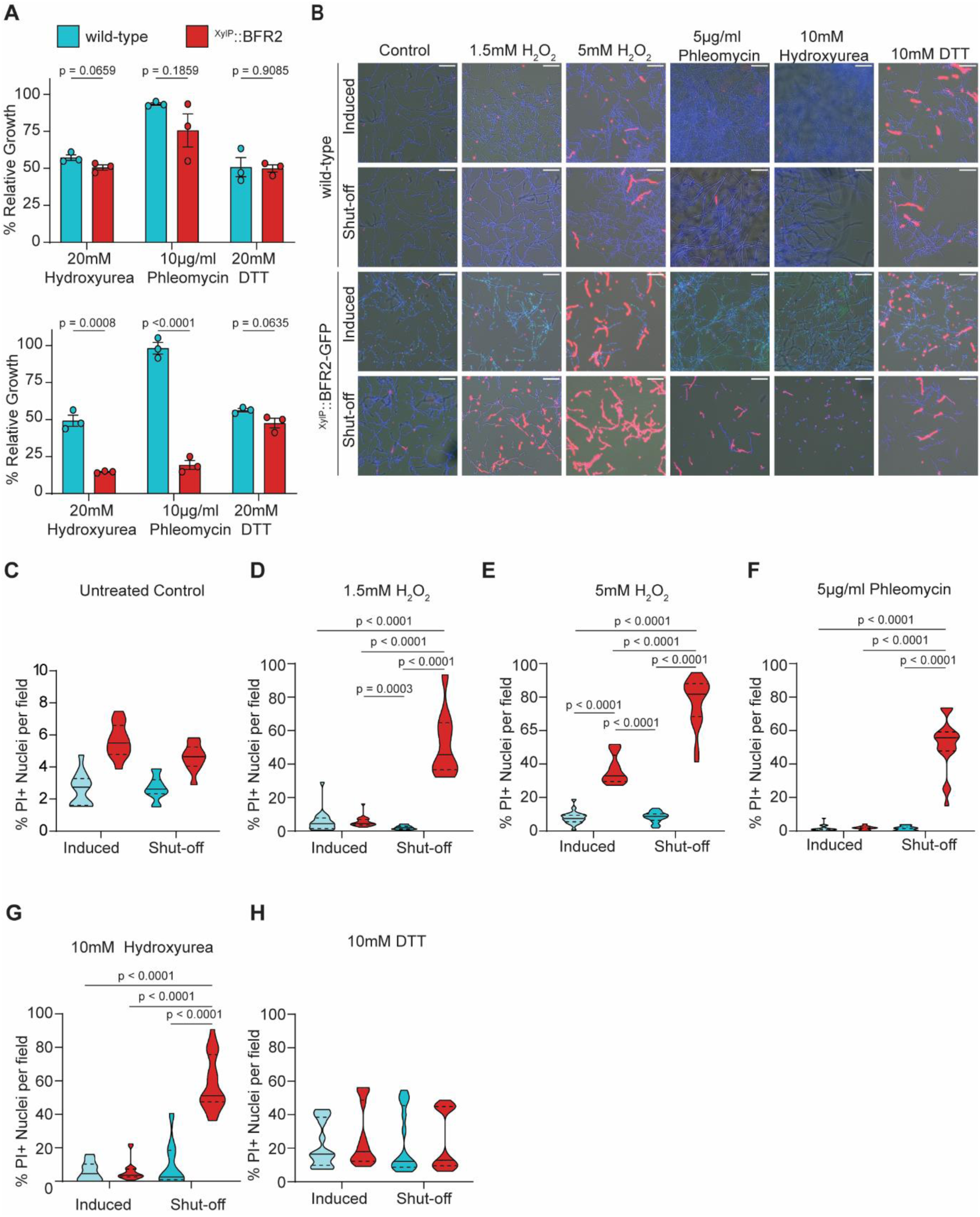
BFR2 is required by *A. fumigatus* for stress tolerance toward oxidative, DNA damage, and replicative stress following germination. (A) Stress challenge on agar plates, determined by radial growth. 1 × 10⁵ resting conidia of wild type and ^XylP::^BFR2 were inoculated onto agar plates containing 1% XMM (top panel) or 0.05% xylose supplemented in 1% GMM (bottom panel), each supplemented with hydroxyurea (20 mM), phleomycin (10 μg/mL), or DTT (20 mM). Plates were incubated at 37°C for 4 days, and radial growth was determined on day 4. Percent relative growth was calculated as radial diameter relative to that on untreated agar plates of the same base medium (1% XMM or 1% GMM with 0.05% xylose, respectively). Data are shown as mean ± SEM of three independent biological replicates, each performed in duplicate. Statistical significance was assessed by t-test for each stress condition and determined at p< 0.05. (B) Fluorescence microscopy of wild type and ^XylP::^BFR2 germlings under stress challenge. BFR2 was pre-induced for 6 hours in 1% XMM. Induced germlings remained in 1% XMM for the entire period, while shut-off germlings were transferred to 1% GMM for an additional 6 hours to deplete BFR2. Germlings were then challenged with hydrogen peroxide (1.5 mM or 5 mM) for 2 hours to induce oxidative stress, phleomycin (5 μg/mL) or hydroxyurea (10 mM), each for 4 hours, to induce DNA damage or replicative stress, respectively, or DTT (10 mM) for 2 hours to induce ER stress. Following challenge, slides were stained with Hoechst to label nuclei. Panels show Hoechst (blue), propidium iodide (red), and BFR2-GFP (green). Non-viable germlings are identified by positive propidium iodide staining. (C–G) Quantification of germling viability after stress challenge, determined as the percentage of Hoechst-positive nuclei that co-stained with propidium iodide. (C) Hydrogen peroxide, 1.5 mM (oxidative stress). (D) Hydrogen peroxide, 5 mM (oxidative stress). (E) Phleomycin, 5 μg/mL (DNA damage). (F) Hydroxyurea, 10 mM (replicative stress). (G) DTT, 10 mM (ER stress). Data are presented as violin plots from two independently performed experiments, with 100 nuclei quantified per condition per strain in each experiment. Statistical significance was assessed by one-way ANOVA with Tukey’s multiple comparisons test Statistical significance was determined at p< 0.05.

Agar plates cannot circumvent the germination constraint imposed by inadequate BFR2 induction at 0.05% xylose, therefore, we employed the two-phase transient induction followed by depletion experimental approach established in the preceding results section to interrogate the stress-responsive role of BFR2 independently of its role in ribosome biogenesis. Wild type and ^XylP::^BFR2 mutant strain germlings were pre-established under full induction conditions for 6h, transferred to either induction or shut-off medium for another 6h, and exposed to a panel host-relevant stressors following the cumulative 12h incubation. Oxidative stress was imposed with hydrogen peroxide at 1.5mM and 5mM; double strand DNA damage with phleomycin at 5 µg/ml; replication stress with hydroxyurea at 10mM; and ER stress with DTT at 10mM. Germling viability was quantified by co-staining nuclei with Hoechst 33342 and propidium iodide (PI), with the proportion of PI-positive nuclei serving as a direct readout of membrane integrity loss and cell death. To assess whether the experimental system itself did not compromise baseline viability, untreated germlings were assessed across all strains and media conditions.. Under both induction and shut-off conditions, PI incorporation remained below 10% in untreated wild type and ^XylP::^BFR2 germlings, although conditional mutants showed slightly higher basal PI staining (5.6% in induced conditions and 4.6% in shut-off conditions) compared to the wild type (2.6% in both induced and shut-off conditions) (Figure 3B,C). Shut-off conditions were not accompanied by a marked increase in PI incorporation establishing that the media switch does not induce stress or alter baseline viability and corroborating the normal physiology of the conditional mutant under both expression states. BFR2 depletion rendered *A. fumigatus* germlings markedly hypersensitive to oxidative stress. Exposure to hydrogen peroxide produced a pronounced increase in PI-positive nuclei specifically in ^XylP::^BFR2 germlings under shut-off conditions (51.9% and 77.25% in 1.5mM and 5mM hydrogen peroxide respectively), relative to both the induced ^XylP::^BFR2 -GFP strain and wild type controls under either media condition (Figure 3B, D, E). This heightened sensitivity was observed at both 1.5 mM and 5 mM hydrogen peroxide, demonstrating a dose-independent requirement for BFR2 in oxidative stress tolerance rather than a threshold effect at a single concentration (Figure 3D, E). BFR2 depletion similarly conferred hypersensitivity to genotoxic and replication stress. Wild type germlings displayed modest, media-independent sensitivity to both agents, with approximately 1.6% and 1.5% killing under phleomycin challenge, 5.9% and 9.6% killing under hydroxyurea across both XMM and GMM conditions respectively (Figure 3B, F, G), confirming that the media switch does not influence genotoxic stress tolerance in a BFR2-replete background. The ^XylP::^BFR2 strain under induction conditions exhibited similar levels of sensitivity relative to wild type, with approximately 1.8% killing under phleomycin and 5.9% under hydroxyurea. Strikingly, BFR2 depletion in GMM dramatically exacerbated this sensitivity to 51.9% during phleomycin challenge and 58.1% during hydroxyurea challenge (Figure 3F, G), demonstrating that BFR2 is required for survival under both DNA damage and replication stress. Notably, these phenotypes emerged despite prior BFR2 induction before the shut-off phase, indicating that pre-existing BFR2 protein was insufficient to confer durable protection once expression was withdrawn (Figure 3F, G). Treatment with 10mM DTT used to induce ER stress resulted in comparable killing across strains and expression conditions. Killing was approximately 19% and 18% in wild type germlings under induced and shut-off conditions respectively, compared to 20% and 18% in ^XylP::^BFR2 germlings. No statistically significant difference was detected between groups, indicating that BFR2 depletion did not significantly alter susceptibility to DTT under the conditions tested (Figure 3H).

The stress-dependent killing phenotypes observed upon BFR2 depletion differed not only in magnitude but in the developmental stage of the fungus most affected. Under replication stress induced by hydroxyurea, PI incorporation was predominantly observed in germlings, indicating that cell death occurred at the earliest stage of polarised growth (Figure 3G). Under DNA damage induced by phleomycin, by contrast, PI-positive nuclei were concentrated in hyphal compartments, demonstrating that BFR2 depletion was lethal preferentially during active hyphal extension rather than during germination. Collectively, these results establish that BFR2 is continuously required for stress tolerance following germination, that prior induction of BFR2 is insufficient to confer durable protection once expression is withdrawn, and that its requirement is both stress-type-specific and stage-specific across fungal development.

### BFR2 is required for *A. fumigatus* to withstand host immune pressure *in vitro* and *in vivo*

To determine whether BFR2 contributes to survival under immune-mediated killing, we adapted the two-phase transient induction followed by depletion protocol for an *in vitro* neutrophil challenge assay. Conidia were induced for 6 hours under full BFR2 induction conditions, followed by a 1-hour period during which the depletion process was initiated in 1% GMM. Calcofluor white (CFW) pre-stained conidia were then co-cultured with primary mouse neutrophils for 6 hours in either fully induced medium (RPMI supplemented with 1% xylose) or shut-off medium (RPMI alone). We determined neutrophil uptake of conidia, conidia viability was assessed by loss of HRFP fluorescence and quantified by colony-forming units (CFU). Neutrophil uptake was comparable across all strains and conditions, with no significant difference detected between wild type and ^XylP::^BFR2 mutant conidia under either induction or shut-off medium. Uptake of wild type conidia was 26.62% under induced conditions and 29.35% under shut-off conditions, while uptake of ^XylP::^BFR2 conidia were 27.18% under induced conditions and 24.63% under shut-off conditions (Figure 4A), establishing that BFR2 depletion does not alter conidial surface recognition or phagocytic uptake by neutrophils. By contrast, BFR2 depletion markedly reduced conidia viability following neutrophil challenge. Wild type conidia maintained similar viability under both conditions, with approximately 75.63% survival under induction and 78.39% under shut-off (Figure 4B). The ^XylP::^BFR2 strain under induction conditions showed a modest, non-significant reduction in viability relative to wild type, with approximately 56.825% survival, whereas BFR2 depletion under shut-off conditions resulted in a severe loss of viability, with survival significantly reduced to approximately 26.5% (Figure 4B). CFU-based quantification corroborated these findings. Wild type survival was approximately 69.59% under induction and 59.11% under shut-off, with no significant difference between conditions. ^XylP::^BFR2 survival under induction was approximately 55.73%, not significantly different from wild type; however, ^XylP::^BFR2 survival under shut-off was reduced to approximately 25.84%, significantly lower than both the induced ^XylP::^BFR2 mutant and the wild type shut-off control (Figure 4C). Taken together, these data demonstrate that BFR2 depletion sensitizes *A. fumigatus* conidia to neutrophil-mediated killing without affecting the initial phagocytic recognition step, and that prior BFR2 induction is insufficient to sustain viability once expression is withdrawn during immune challenge.

**Figure 4.**
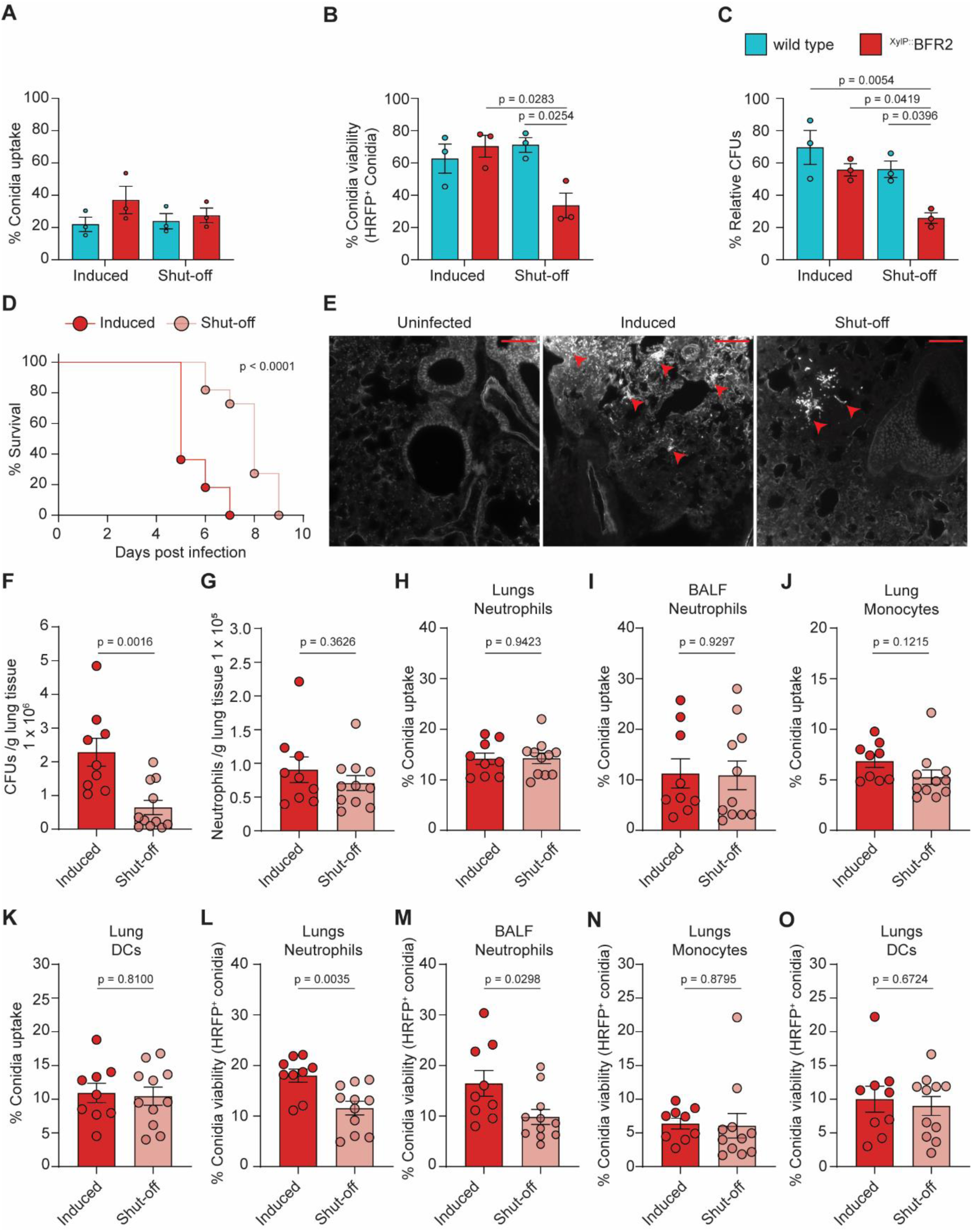
Transient induction followed by depletion of BFR2 compromises conidial viability following phagocytosis by bone marrow–derived neutrophils in vitro and lung and bronchoalveolar lavage fluid (BALF) neutrophils *in vivo*. (A–C) In vitro neutrophil challenge. Bone marrow–derived neutrophils were harvested from wild type C57BL/6J mice and co-incubated with swollen conidia of the HRFP and ^XylP::^BFR2-HRFP strains. BFR2 was pre-induced in 1% XMM for 6 hours, followed by a 1-hour incubation in 1% GMM to initiate depletion. Conidia were then labelled with calcofluor white (100 μg/mL) and co-incubated with neutrophils (1 × 10⁶) at a multiplicity of infection (MOI) of 1:1. A subinhibitory concentration of voriconazole (0.5 μg/mL) was included to prevent conidial germination, and IFN-γ (1 ng/mL) was added to induce phagocytosis by bone marrow– derived neutrophils. Data are shown as mean ± SEM of three independent experiments performed in triplicate. Statistical significance was assessed by one-way ANOVA with Tukey’s multiple comparisons test (p-value <0.05); comparisons without a p value indicated were not significant. (A) Conidial uptake by bone marrow–derived neutrophils. (B) Conidial viability after neutrophil challenge, indicated by the proportion of HRFP-positive conidia. (C) Conidial viability after neutrophil challenge, presented as % relative CFU. **(D–I) *In vivo* infection.** (D) Survival following infection of C57BL/6J mice immunosuppressed with Cyclophosphamide 150mg/ml 3 days before infection, 100mg/ml one day before infection, and then every two days. Significance was assessed by Mantel-Cos log-rank test. Mice were euthanized on reaching humane end point defined by loss of 20% of initial body weight. (E) Representative lung section histology slides stained with calcofluor white (CFW). Images shown are uninfected, induced, and shut off groups at 48h post infection. Red arrowheads indicate A. fumigatus hyphae representing foci of invasive growth. Scale bar, 80um. Images are representative of 3 mice per group.(F-O) Immunocompetent C57BL/6J mice were challenged with 30 million ^XylP::^BFR2-HRFP conidia. BFR2 was pre-induced in 1% XMM for 6 hours, followed by a 1-hour incubation in 1% GMM to initiate depletion. Conidia were subsequently labelled using the FLARE (fluorescent Aspergillus reporter) for identification *in vivo*. Mice were administered 10% xylose in drinking water 24 hours prior to infection. Infection was carried out for 24 hours, after which lung fungal burden (CFUs), neutrophil recruitment, conidial uptake, conidial killing by lung and BALF neutrophils, monocytes and dendritic cells were assessed.. (F) Fungal burden expressed as CFUs of gram of lung tissue. (G) Neutrophil recruitment to the lungs after 24-hour infection, presented as neutrophils per gram of lung tissue. (H-K) Conidial uptake by lung neutrophils (H) and BALF neutrophils (I), lung monocytes (J), lung dendritic cells (DCs, K). (L-O) Conidia viability after in vivo challenge, expressed as % HRFP-positive conidia, in lung neutrophils (L), BALF neutrophils (M), lung monocytes (N), and lung DCs (O). Data are in F-O are shown as mean ± SEM of 9 mice (induced) and 11 mice (shut-off). Statistical significance was assessed by unpaired student’s t-test and determined at p <0.05.

To determine whether BFR2’s role extends to virulence in *A. fumigatus*, we extended the transient induction/shut-off strategy to an immunosuppressed murine model of invasive aspergillosis. Conidia of ^XylP::^BFR2 mutant were pre-swollen in XMM for 6h and either maintained under inducing conditions or transferred into GMM to initiate the depletion of BFR2 in the shut-off condition for 1h prior to infection, and the mice were challenged with 5 × 10^4^ conidia. To sustain induction in vivo, mice infected with induced conidia received 10% xylose in their drinking water 24h before infection, and ad libitum throughout the experiment. This regimen has been previously shown to be sufficient for xylose-driven gene expression in mice.

Survival was monitored for 9 days post-infection. Mice infected with induced conidia succumbed more rapidly reaching 100% mortality by day 7, whereas mortality in the shut-off group was delayed, reaching 100% by day 9 (Figure 4D). To assess early tissue invasion, lungs were collected 48h post infection and fungal elements were visualized by calcofluor white staining of tissue sections. Lungs from mice infected under inducing conditions showed extensive hyphal growth and germlings occupying large areas of lung tissue, whereas the shut-off group showed fewer and smaller foci of fungal growth (Figure 4E).

Together, these data indicate that BFR2 is not essential for virulence in the strict sense, but contributes to pathogenicity, and that depleting BFR2 at the point of infection delays tissue invasion and extends survival. The presence of germinating conidia in the shut-off group indicates that BFR2-depleted conidia retain their capacity to grow in an immunosuppressed lung, albeit more slowly, which accounts for delayed rather than abolished mortality.

To determine whether the secondary role of *A. fumigatus* BFR2 in stress response extends beyond *in vitro* beyond the neutrophil challenge, wild type C57BL/6J mice were challenged with 30 million pre-swollen ^XylP::^BFR2 conidia. Pre-induction of BFR2 was performed in 1% XMM for 6 hours, followed by a 1-hour period in 1% GMM to initiate BFR2 depletion prior to infection, mirroring the two-phase induction/shut-off protocol used above. Prior to infection, mice were administered 10% xylose as above. Mice were challenged for 24 hours, after which lung fungal burden and neutrophil recruitment into the lungs was determined. Fungal burden in the lungs, determined by CFU quantification, was significantly reduced under shut-off conditions, with an average of 2.28 × 10⁶ CFU per gram of lung tissue under induced conditions compared to 0.64 × 10⁶ CFU per gram of lung tissue under shut-off conditions (Figure 4F). Neutrophil recruitment into the lungs did not differ significantly between induced and shut-off conditions (Figure 4G). Recruitment reached 0.9 × 10⁵ cells per gram of lung tissue under induced conditions and 0.7 × 10⁵ cells per gram of lung tissue under shut-off conditions.

Conidial uptake by recruited lung and BALF neutrophils, lung monocytes and dendritic cells were unaffected by BFR2 shut-off (Figure 4H-K). In lung neutrophils, conidial uptake was 14.15% under induced conditions and 14.26% under shut-off conditions (Figure 4H). In BALF neutrophils, conidial uptake was 11.25% under induced conditions and 10.88% under shut-off conditions (Figure 4I). Similar trends of conidia uptake were observed in lung monocytes and dendritic cells with uptake of 6.84% under induced conditions and 5.27% under shut-off conditions in lung monocytes (Figure 4J) and 10.93% under induced conditions and 10.4% under shut-off conditions in lung dendritic cells (Figure 4K). Thus, the depletion for BFR2 expression did not affect the efficiency with which recruited phagocytes and lung leucocytes take up conidia *in vivo*.

Although neutrophil recruitment and phagocytic uptake were unaffected by BFR2 depletion, conidial viability was reduced in both lung and BALF neutrophils following shut-off. Conidial viability was assessed using Histone 2A-mRFP fluorescence, with loss of RFP fluorescence indicating non-viable conidia. In lung neutrophils, 17.99% of conidia remained viable under induced conditions, compared to 11.56% under shut-off conditions. A comparable reduction was observed in BALF neutrophils, with 16.45% of conidia viable under induced conditions compared to 9.81% under shut-off conditions (Figure 4L,M). On the other hand, conidia viability within lung monocytes (6.4% of conidia viable under induced conditions and 5.86% under shut-off conditions) and dendritic cells (9.99% of conidia viable under induced conditions and 8.99% under shut-off conditions) were unchanged (Figure 4N-O).

Taken together, these data indicate that while BFR2 depletion did not impair neutrophil recruitment or phagocytic uptake of conidia by neutrophils, lung monocytes and dendritic cells *in vivo,* it significantly compromised conidia viability specifically within neutrophils and resulted in an overall fungal burden in the lungs.

## Discussion

Currently approved antifungal drugs have concentrated on the fungal cell wall and membrane, leaving fungal stress networks which the fungi depend on to survive within the host largely unexplored as therapeutic targets. Here we show that BFR2 in *A. fumigatus* performs a role beyond its canonical function in ribosomal biogenesis. We showed that it is required for survival of oxidative, DNA damage and replicative stress, and for fungal survival during neutrophil challenge and *in vivo* infection.

In fungi, especially *A. fumigatus*, ribosomal biogenesis factors have been characterized by their canonical ribosomal function role, and sometimes in relation to environmental stressors^27,54,55^. Their capacity for additional regulatory functions, which is a well-established role for nucleolar proteins in mammalian systems has remained largely unexamined in fungi. In *A. fumigatus*, the clearest precedence of stress-linked ribosome biogenesis factor is CgrA where its disruption reduced the ability of the fungus to accelerate growth at 37°C and reduced virulence in an immunosuppressed mouse model^27,54^. This phenotype was traced back to alterations in ribosomal subunit ration 60S:40S and the accumulations of half-mer ribosomes. It was proposed that fungal nucleolus might parallel its mammalian counterpart in coordinating stress responses and DNA repair alongside ribosomal biogenesis, however, CgrA has remained the only ribosomal biogenesis factor in *A. fumigatus* that has been experimentally linked to a defined stress phenotype. Our findings extend this coupling of ribosomal biogenesis to other stress phenotypes by showing that showed that BFR2 couples ribosome biogenesis to oxidative, genotoxic and replicative stress resistance, and to survival within the host. This indicates a functional expansion that is not limited only to mammalian biology but extends to major fungal pathogens.

Consistent with studies in *Saccharomyces cerevisiae* and mammalian cells, BFR2 proved essential for 18S rRNA biogenesis, germination and vegetative growth, confirming its conserved role in ribosome biogenesis (Figure 1). Critically, however, acute depletion of BFR2 once ribosome assembly was already established selectively impaired resistance to oxidative, replication and genotoxic stress, without affecting ribosomal RNA abundance, global translation or biomass accumulation over the same experimental period (Figure 2-3). These observations raise the possibility that BFR2’s canonical role in ribosome biogenesis extends to a role in stress adaptation, consistent with a model in which fungal ribosomal biogenesis proteins are required for responses to stress.

The emerging view of the nucleolus as a signaling center provides an intriguing evolutionary context for interpreting these functions. In mammalian cells, the BFR2 homolog AATF (Che-1) has evolved beyond its canonical role in ribosome biogenesis to coordinate DNA damage responses, replication stress signaling, nucleolar homeostasis and cell survival, largely through p53-dependent pathways^45,49^. Fungi lacks p53 and its associated regulatory network^56^, yet BFR2 in *A. fumigatus* performs a comparable stress-coordinating role, raising the possibility that this function within the BFR2/AATF family predates the metazoan-specific signaling architecture built around it. Under this model, the ancestral role of BFR2/AATF extended beyond ribosome biogenesis to encompass coordination of the cellular stress response, with p53-dependent regulation representing a later evolutionary elaboration in animals rather than the origin of the function itself.

A central finding of this study is that BFR2 depletion does not produce a uniform stress-sensitive phenotype but instead selectively compromises tolerance to oxidative, replication and genotoxic stress while leaving the response to endoplasmic reticulum stress largely unaffected (Figure 3). This pattern is particularly informative because it argues against a simple model in which all phenotypes arise as secondary consequences of impaired ribosome biogenesis or reduced translational capacity. Ribosome production underpins virtually every aspect of cellular physiology, and therefore a generalized defect in ribosome assembly would be expected to broadly impair adaptation to mechanistically distinct stresses. Instead, the preservation of DTT tolerance, despite pronounced hypersensitivity to hydrogen peroxide, hydroxyurea and phleomycin, indicates that BFR2 is unlikely to function solely as a constitutive regulator of protein synthesis or general cellular fitness.

Rather, the selective nature of the phenotype suggests that BFR2 contributes to a more defined stress adaptation programme. Oxidative stress, replication stress and DNA damage are mechanistically distinct insults, yet all converge on the maintenance of genome integrity. Reactive oxygen species generate oxidative DNA lesions, hydroxyurea induces replication fork stalling through depletion of intracellular deoxyribonucleotide pools, and phleomycin directly produces DNA double-strand breaks. The shared requirement for BFR2 under these conditions therefore raises the possibility that its stress-associated function is linked to pathways that preserve genome stability during environmental challenge. This logic has precedent within the genus: in *Aspergillus nidulans*, loss of the ATR homolog UvsB confers sensitivity to both hydroxyurea and DNA-damaging agents, indicating that a single checkpoint node can govern resistance across mechanistically distinct genotoxic and replicative insults^57^. A comparable precedent exists for the BFR2 family itself. The depletion of the mammalian ortholog AATF increases DNA damage and reduces genotoxic tolerance through direct recruitment to damage sites and interaction with the repair protein PARP1, independently of its role in ribosome biogenesis^58–60^. Although the molecular basis of this role in *A. fumigatus* remains unresolved, our findings narrow the spectrum of plausible mechanisms and argue against a model of generalized proteostatic failure (Figure 2I).

The lack of sensitivity to DTT provides an important internal control for this interpretation (Figure 3B). DTT disrupts disulphide bond formation within the endoplasmic reticulum, activating the unfolded protein response and placing substantial demands on protein folding and proteostasis rather than directly challenging genome integrity. If the stress phenotypes observed following BFR2 depletion simply reflected insufficient ribosome production or reduced global translation, sensitivity would be expected across all stress conditions, including ER stress. Instead, the preservation of DTT tolerance suggests that core translational capacity remains sufficient to support adaptation to at least some environmental insults and that the stress-associated function of BFR2 is selective rather than universal. This distinction argues that BFR2 is unlikely to act merely as a housekeeping ribosome biogenesis factor and instead supports a model in which it participates in pathways specifically associated with oxidative damage, replication stress or DNA damage responses. This mode of coupling is also mechanistically distinct from the well-characterised RP–MDM2–p53 surveillance axis, in which genome-integrity signalling is triggered by ribosome biogenesis impairment itself ^40,61^. In our system, rRNA abundance, translation and biomass accumulation remained intact throughout the depletion window, indicating that BFR2’s contribution to stress resistance does not depend on, and is not a downstream consequence of, a detectable ribosome biogenesis defect (Figure 2).

The physiological relevance of the stress-associated functions of BFR2 is underscored by its requirement for survival during neutrophil-mediated killing. Neutrophils constitute the first line of innate immune defence against *A. fumigatus* and are indispensable for preventing invasive aspergillosis, as evidenced by the heightened susceptibility of both NADPH oxidase-deficient humans and mice to invasive disease^20,62–64^. Following phagocytosis, conidia are exposed to a complex antimicrobial environment: intracellular killing requires the fusion of granules containing cathepsin and neutrophil elastase together with activation of the NADPH oxidase, generating reactive oxygen species (ROS) and hypochlorous acid (HOCl) alongside antimicrobial proteases, while neutrophils additionally sequester essential nutrients through nutritional immunity. Many of the intracellular antimicrobial mechanisms deployed against conidia either directly induce DNA damage or generate oxidative lesions that compromise genome integrity, necessitating robust stress adaptation and repair pathways for fungal survival.

Against this biological backdrop, the marked hypersensitivity of BFR2-depleted swollen conidia to neutrophil-mediated killing is particularly informative (Figure 4B-C). Importantly, BFR2 depletion did not alter neutrophil uptake, indicating that the increased susceptibility cannot be attributed to altered conidial recognition, cell wall architecture or phagocytosis (Figure 4A). Instead, the defect became apparent only after internalisation, demonstrating that BFR2 is required for survival within the hostile intracellular environment generated by activated neutrophils. The concordance between the selective stress phenotypes observed *in vitro* and the immune-cell killing assay therefore argues that the neutrophil phenotype is not an independent observation but rather a biological manifestation of the stress adaptation defects identified throughout this study.

The close correspondence between the stress assays and the neutrophil challenge further strengthens this interpretation. Acute depletion of BFR2 selectively impaired resistance to oxidative stress, replication stress and DNA damage, all of which are encountered directly or indirectly during neutrophil attack. Reactive oxygen species generated during the oxidative burst not only damage proteins and lipids but also produce oxidative DNA lesions and replication-associated stress, while secondary oxidants such as HOCl amplify these effects. Consequently, the susceptibility of BFR2-depleted fungi to neutrophil-mediated killing is consistent with an inability to withstand the combined oxidative and genotoxic pressures imposed by the host innate immune response rather than a defect in immune recognition or fungal uptake.

This translatability of the BFR2 phenotype into immune-cell susceptibility is notable against the broader pattern reported for oxidative-stress-defense genes in *A. fumigatus*, where increased sensitivity to reactive oxygen species in vitro has generally failed to predict a corresponding increase in phagocyte-killing susceptibility or virulence. Deletion of the conidial catalase *catA* increases sensitivity to hydrogen peroxide without altering conidial killing by alveolar macrophages or virulence in vivo, and combined loss of the mycelial catalases *cat1* and *cat2* similarly fails to increase susceptibility to killing by human neutrophils despite abolishing catalase activity entirely. Deletion of the fungal superoxide dismutase increases susceptibility to macrophage killing without producing a corresponding difference in virulence in animal infection models. This pattern is generally attributed to redundancy among the multiple, overlapping systems *A. fumigatus* deploys against reactive oxygen species; the peroxiredoxin Aspf3 is one of the few characterised exceptions in which oxidative-stress sensitivity and reduced fitness within the host concur. Positioned against this backdrop, the consistent translation of the BFR2 stress phenotype into neutrophil-killing susceptibility is atypical and suggests that BFR2 does not function as one redundant node within a single detoxification pathway but instead occupies a more integrative position spanning the oxidative, replicative and genotoxic stress responses examined in this study.

These findings reinforce the emerging concept that ribosome biogenesis factors can influence host-pathogen interactions through functions extending beyond cellular growth. While BFR2 is essential for ribosome biogenesis, our transient depletion strategy demonstrates that stress-associated defects arise under conditions in which ribosomal RNA abundance, global translation and biomass accumulation remain largely intact. The enhanced neutrophil killing observed under these conditions therefore supports a model in which BFR2 contributes directly to host-associated stress adaptation, linking nucleolar function to fungal survival during innate immune attack. More broadly, these observations suggest that multifunctional nucleolar proteins may represent attractive antifungal targets, as their inhibition could simultaneously compromise fungal growth and diminish the capacity of the pathogen to resist immune-mediated stress.

## Methods

### Fungal Strains and Growth Conditions

Mutant strains were generated in *A. fumigatus* wild type from the American Type Culture Collection (ATCC) 46645 background and in an ATCC histone 2A::monomeric red fluorescent protein (HRFP) reporter strain^26,65^. The HRFP strain constitutively expresses a Histone 2A-RFP fusion protein, which serves as a fluorescent viability marker in which the loss of HRFP fluorescence indicate non-viable cells. Wild type strains were maintained on 1% glucose minimal media (1% GMM), while mutant strains were maintained on 1% xylose minimal media (1% XMM). All strains were stored as conidial stocks at −80C in 25% glycerol. 1% GMM consisted of of 6 g/L NaNO₃, 0.52 g/L KCl, 0.52 g/L MgSO₄·7H₂O, 1.52 g/L KH₂PO₄, 2.2 mg/L ZnSO₄·7H₂O, 1.1 mg/L H₃BO₃, 0.5 mg/L MnCl₂·4H₂O, 0.5 mg/L FeSO₄·7H₂O, 0.16 mg/L CoCl₂·5H₂O, 0.16 mg/L CuSO₄·5H₂O, 0.11 mg/L (NH₄)₆Mo₇O₂₄·4H₂O, 5 mg/L Na₄EDTA, and 10 g/L glucose, with pH adjusted to 6.5. 1% XMM was identical to 1% GMM except that glucose was replaced with 10g/L xlyose. Solid agar media were prepared by adding 1.5% agar to the above formulations. All media were autoclaved prior to use. Liquid media used for fluorescence microscopy were filter sterilized through 0.2μm filter to generate optically clear media.

### Mutant Strain Generation

The endogenous *BFR2* locus was targeted by CRISPR-Cas9-mediated cleavage using a single guide RNA directed to a site near the translational start codon. Repair of the resulting double-strand break by non-homologous end joining generated an indel that introduced a premature stop codon near the start of the coding sequence, confirmed by amplicon sequencing (Plasmidsaurus). The 3xFlag-BFR2-GFP complementation construct, under the control of the xylose-inducible promoter, was introduced into this background by microhomology mediated CRISPR-Cas9 and integrated ectopically at a genomic locus that was not mapped (Figure S1B). Successful integration was confirmed by PCR amplification of the full-length construct, detection of GFP expression, and phenotypic complementation.

### Radial growth assay

Radial growth was assessed by spot inoculation of 1 × 10⁵ resting conidia, in a 5 µl volume, onto the center of solid agar plates prepared with fully inducing 1% xylose minimal medium (XMM), fully repressing 1% glucose minimal medium (GMM), or 1% GMM supplemented with 1%, 0.05%, or 0.04% xylose. Plates were incubated at 37°C for 4 days.

### Germination assay

Germination rate was determined using conidia harvested from 4-day-old XMM plates. Conidia were counted and adjusted to 5 × 10⁵ conidia/ml in 1% GMM (fully repressing), 1% XMM (fully inducing), or 1% GMM supplemented with 1%, 0.5%, 0.3%, 0.1%, 0.05%, or 0.02% xylose. Conidial suspensions were inoculated into an 8-well chambered microscopy slide (Ibidi) and incubated at 37°C for 10 hours, at which point germination was scored. Images were acquired at 10 hours using an inverted motorized Nikon Eclipse Ti2 fluorescence microscope.

### RNA extraction and RT-qPCR

Mycelia from liquid shaking cultures were flash-frozen in liquid nitrogen, lyophilized, and ground with 0.5 mm silica beads using a Precellys homogenizer (1 cycle, 30 s). Ground mycelia were homogenized in Trizol at a ratio of 50 mg mycelia per 1 ml Trizol. Chloroform (200 μl) was added, and the aqueous phase was collected following centrifugation. RNA (10 µg) was DNase-treated using the TURBO DNA-free kit (Invitrogen) according to the manufacturer’s instructions, and cDNA was synthesized using the High-Capacity cDNA Reverse Transcription kit (Applied Biosystems).

RT-qPCR was performed using Fast SYBR Green Master Mix (Applied Biosystems, 4385612), with data collected on a StepOnePlus Real-Time PCR System (Applied Biosystems). Primer pair concentrations were optimized prior to generating amplification efficiency curves; only primers with amplification efficiency between 95–105% were used. Non-template controls were included in every run to confirm the absence of contaminating amplification and melt curve analysis was performed following each run to confirm the absence of non-specific amplification products. Fold change in BFR2 transcript abundance was calculated using the 2^−ΔΔCt^ method, normalized to the *A. fumigatus* actin gene (actA). FLAG and GFP, used as surrogates to assess BFR2 expression under the xylose-inducible promoter, were instead quantified as absolute 2^−ΔCt^ values normalized to actA expression, since the wild-type strain lacks these sequences and cannot serve as a ΔΔCt calibrator.

### Mycelial Biomass Accumulation Assay

Biomass accumulation was assessed by inoculating 5 × 10⁷ conidia in 1% GMM (fully repressing) and 1% XMM (fully inducing) conditions, with shaking at 150 rpm at 37°C for 24 hours. Mycelium was harvested, flash-frozen in liquid nitrogen, and lyophilized for 4 days. Lyophilized mycelium was weighed using an analytical balance to determine biomass. To assess biomass accumulation under an induction shut-off regime, conidia (5 × 10⁷) were incubated in 1% XMM to pre-induce BFR2 expression for 3 or 6 hours, washed with PBS-Tween, and then shifted to 1% GMM to initiate BFR2 depletion for the remainder of a 24-hour incubation period at 37°C with shaking at 150 rpm. Mycelium was then harvested and processed as described above.

### Stress induced conidia/germlings killing assays

Conidia (5 × 10⁵) were immobilized on removable microscope chambered slides pre-coated with poly-L-lysine and incubated for 6 hours in 1% XMM to pre-induce BFR2 expression in swollen conidia. For the shut-off condition, immobilized conidia were subsequently washed with PBS and incubated in 1% GMM for a further 6 hours to initiate BFR2 depletion. Stressors used in this assay include 1.5 mM or 5 mM hydrogen peroxide (tested as separate conditions) to induce oxidative stress, 5 µg/ml phleomycin to induce DNA damage stress, 10 mM hydroxyurea to induce replication stress, and 10 mM DTT to induce ER stress — were added to the cells and incubated for 2 hours (oxidative stress), 4 hours (DNA damage and replication stress), or 2 hours (ER stress). Following the appropriate incubation period, cells were stained with Hoechst 33342 and propidium iodide (PI) for 30 minutes at 37°C and imaged using an inverted motorized Nikon Eclipse Ti2 fluorescence microscope. Images were acquired on the DAPI channel (Hoechst), RFP channel (PI), and GFP channel (BFR2-GFP), and analyzed using ImageJ. For each slide, Hoechst-stained nuclei were quantified across 100 germlings per condition, and PI-stained germlings were quantified for the same population. Germling viability was determined as the percentage of PI-positive nuclei.

### Mice

Male and female C57BL/6J mice, 8-10 weeks of age, were used for all experiments. Mice were bred in-house under specific pathogen-free conditions at the animal facility of the Hebrew University of Jerusalem. All experiments were performed in accordance with institutional standards for the care and use of laboratory animals. The facility was maintained under standardized environmental conditions, with room temperature held at 22 ± 2°C and relative humidity controlled between 30-50%. Animals were housed under a 12 h light/dark cycle. All procedures were approved by the Hebrew University of Jerusalem Institutional Animal Care and Use Committee (IACUC protocol HU-24-17584-5).

### *In vitro* neutrophil phagocytosis assay

Conidial induction and shut-off were performed as previously described, with slight modifications. Swollen conidia were pre-induced for 6 h in 1% XMM at 37°C with shaking at 150 rpm to induce BFR2 expression. Conidia were then washed with 0.0025% PBS-Tween and incubated in 1% GMM at 37°C for 1 h to initiate BFR2 depletion in the shut-off condition. Conidia in the induction condition remained in XMM for the equivalent period. Following this step, conidia were washed in PBS-Tween and labeled with 100 µg/mL Calcofluor White for 20 min at room temperature, washed, and resuspended in PBS.

Bone marrow-derived neutrophils were isolated from femurs and tibias of C57BL/6J mice. Mice were anesthetized with sodium pentobarbital (5 mg/mL) prior to bone harvest. An incision was made along the midline and around both rear legs, and the skin was removed from the legs. Legs were severed above the hip joint, and femurs and tibias were isolated by trimming away muscle and connective tissue and separated by severing the patellar tendon and gently bending the knee joint backward. Cleaned bones were placed in cold PBS on ice.

Under sterile conditions, bones were sterilized in 70% ethanol for 3 min. Bone ends were cut, and marrow was flushed with cold PBS into a Petri dish and collected into a 50 mL conical tube, with each bone washed with at least 2 mL of PBS. The suspension was centrifuged at 1,500 rpm for 5 min, and the supernatant was removed. The pellet was resuspended in 1 mL of 1X RBC lysis buffer (sufficient for bone marrow from four mouse legs) for 5-10 min at room temperature, followed by addition of 9 mL of PBS to neutralize the lysis reaction. The suspension was filtered through a 100 µm strainer, centrifuged again at 1,500 rpm for 5 min, and the supernatant was removed. Cells were resuspended in 10 mL of RPMI/PBS, counted using a haemocytometer, and adjusted to 1 × 10^6^ cells/ml.

Bone marrow-derived neutrophils were separated into RPMI medium alone or RPMI supplemented with 1% XMM, corresponding to the shut-off and induction conditions, respectively. Labelled conidia were resuspended in the corresponding medium for each condition, matching their prior processing.

Neutrophils and conidia were co-incubated at a multiplicity of infection (MOI) of 1, with a final concentration of 1 × 10^6^/mL for each, in the presence of voriconazole (0.5 µg/mL) and IFN-γ (1 ng/mL) to activate bone marrow-derived neutrophils. Sub-inhibitory concentrations of voriconazole (0.5 µg/mL) were added to prevent germination. Conidia-only and neutrophil-only wells were included as controls. Co-incubation was carried out for 6h at 37°C with 5% CO2.

Following co-incubation, cell suspensions were centrifuged, and the supernatant was removed. Cells were incubated in staining solution for 20 min on ice and subsequently stained with CD45.2 (PerCP-Cy5.5) and Ly6G (AF700) antibodies for 20 minutes. Stained cell suspensions were washed, centrifuged, and resuspended in PBS. Samples were analyzed by flow cytometry to assess conidial uptake and neutrophil-mediated killing. Co-incubated cell suspensions were serially diluted and plated on 1% XMM agar for CFU enumeration.

### Murine models of aspergillosis

To assess conidial recruitment into the lungs, phagocytosis and killing by lung and BALF neutrophils, and fungal burden, immunocompetent C57BL/6J mice were used. Mice in the induction group received 10% xylose in their drinking water beginning 24 h before infection, while mice in the shut-off group received untreated drinking water.

Conidial induction and shut-off were performed as previously described, with slight modifications. Swollen conidia were pre-induced for 6 h in 1% XMM at 37°C with shaking at 150 rpm to induce BFR2 expression. Conidia were then washed with 0.0025% PBS-Tween and incubated in 1% GMM at 37°C for 1 h to initiate BFR2 depletion in the shut-off condition, while conidia in the induction condition remained in XMM for the equivalent period. Conidia were subsequently labelled with FLARE as previously described in ^66^. Mice were infected intranasally with 30 × 10^6^ conidia in 50 µL of PBS and were maintained for 24 h following inoculation, at which point lungs and BALF were collected for downstream analysis.

Mice were euthanized by intraperitoneal injection of sodium pentobarbital (100 mg/kg; 2 mg per 20 g mouse, administered as 400 µL of a 5 mg/mL working solution). The trachea, heart, and lungs were exposed, and the hepatic artery was cut to drain systemic circulation. Pulmonary circulation was flushed with 5 mL of PBS via the right ventricle. BALF was collected into a 15 mL tube, and lungs were collected into a separate 15 mL tube containing 5 mL of 5% FBS in PBS. Bone marrow was additionally collected from the legs of one mouse per experiment for use as a compensation control.

Lungs were digested in a solution of collagenase type IV (2.2 mg/mL) and DNase I (100 µg/mL) in 5% FBS/PBS prior to dissociation. Lungs were transferred into GentleMACS C-tubes and processed using a GentleMACS tissue Dissociator, with a 40 min incubation at 37°C when digestion enzymes were used, followed by dissociation on the Lung #2 program. Samples were centrifuged at 1,500 rpm for 5 min, and the supernatant was discarded. Pellets were resuspended in 1 mL of 1X RBC lysis buffer for 5 min at room temperature, and the reaction was neutralized with 9 mL of 5% FBS/PBS. The suspension was filtered through a 40µm filter into a 50 mL conical tube, centrifuged at 1,500 rpm for 5 min, and resuspended in 2 mL of 5% FBS/PBS.

BALF samples were centrifuged at 1,500 rpm for 5 min, and the supernatant was removed, leaving approximately 100 µL (supernatant was retained separately for cytokine analysis where applicable). BALF pellets were resuspended in 300 µL of 5% FBS/PBS, with 200 µL allocated for flow cytometry and 100 µL allocated for RNA and CFU analysis. RBC lysis was not required for BALF samples.

Lung and BALF cell suspensions were transferred to a 96-well microplate, with 200 µL used per lung sample and the entire BALF sample used per well. Samples were centrifuged and resuspended in 50 µL of PBS. Fc block was applied to lung samples only, and cells were stained for flow cytometric analysis. Lung and BALF neutrophils were identified as CD45^+^ (PerCPCy5.5), Ly6C^lo^ (PE Cy7), Ly6G^+^ (AF700) cells using antibodies obtained from BioLegend and diluted according to the manufacturer’s recommendations. Flow cytometry data were acquired on a NovoCyte Quanteon flow cytometer (Agilent) and analyzed using NovoExpress software (v1.6.1, Agilent). Perfused lungs were homogenized in 1 mL of PBS-Tween, serially diluted, and plated to determine colony-forming units (CFU). For histology and survival experiments, mice were immunosuppressed by intraperitoneal injection of cyclophosphamide (Western Medical Supply; 150mg/kg) administered 72 h and 24 h before fungal inoculation, and every 2 days thereafter for the duration of the experiment. For histological analysis, perfused lungs were fixed in 10% neutral-buffered formalin, embedded in paraffin and sectioned at 4 µm. Histological samples were deparaffinized and stained with Calcofluor White (MP Biomedicals).

### Puromycin Incorporation Assay for Bulk Protein Synthesis Measurement

*A. fumigatus* HRFP and ^XylP::^BFR2-HRFP conidia were inoculated into conical flasks containing 50ml 1% Xylose minimal media at a final concentration of 5 × 10^7^ conidia/mL and incubated at 37C with shaking at 150rpm for 6h to generate swollen conidia. For glucose induced shut-off conditions, swollen conidia were harvested and washed with 1x PBS and then incubated in fresh 50ml glucose minimal media to initiate the transcriptional shut-off of the xylose-inducible promoter controlling BFR2 expression. For strains in shut-off conditions, they were maintained in 1% XMM. All incubations were carried out for 6 more hours. As a negative control for translation-dependent puromycin incorporation, cycloheximide (CHX) was added to a control flask containing ATCC HRFP or XylP::BFR2-HRFP grown in XMM at a final concentration of 100 μg/mL, 30 mins prior to the addition of puromycin with continued incubation at 37C, 150rpm. Puromycin was added directly to each culture flask, including the CHX-treated control at a final concentration of 15 μg/mL, and cultures were incubated for an additional 30 min at 37C with shaking at 150rpm to allow incorporation of puromycin into nascent polypeptide chains. An additional culture receiving an equivalent volume of PBS in place of puromycin served as a labelling-negative control. Labelling was halted by immediate transfer of cultures on ice. Mycelial biomass was harvested by filtration on Whatman paper and washed with ice-cold PBS. Mycelial biomass was flash frozen in liquid nit4rogen and grounded in mortar and pestle into fine powder. Powdered mycelium was transferred into 2ml screw cap tubes containing 200uL of 0.5mm zirconia beads and lysed by beat-beating in 400uL of protein lysis buffer (50 mM HEPES (pH 7.4), 150–250 mM NaCl, 1% NP-40, 1 mM EDTA, 1 mM PMSF, 1 mM NaF, and 1 mM NaVO₃, supplemented with 1X protease inhibitor cocktail (Apex Bio)). Lysates were clarified by centrifugation at 20,800 x g for 20 min at 4C, and the resulting supernatants were transferred to fresh, pre-chilled tubes. Protein concentration was determined using Bradford assay, and samples were normalized to equal total protein concentrations. 30ug proteins were loaded per lane for western blot in 4x Laemmli sample buffer containing DTT and denatured by boiling at 95C for 5 mins. Denatured protein samples were resolved on 12% SDS-PAGE and transferred to membranes for immunoblotting. Membranes were blocked in 5% skim milk in 1% tween-TBS buffer for 1h at room temperature with rocking, followed by washing. Puromycin-labelled polypeptides were detected by incubation with anti-puromycin monoclonal antibody (clone 12D10) diluted 1:30000 in 5% BSA/TBST overnight at 4C, followed by washing and incubation with an HRP-conjugated anti-mouse secondary antibody diluted 1:10000 in 5% skim milk/TBST for 1h at room temperature. Membranes were washed and developed by chemiluminescence.

