## Supplementary material for "Ribosomal biogenesis factor links ribosome homeostasis to host relevant stress survival in *Aspergillus fumigatus*": Figure S1, Figure S2

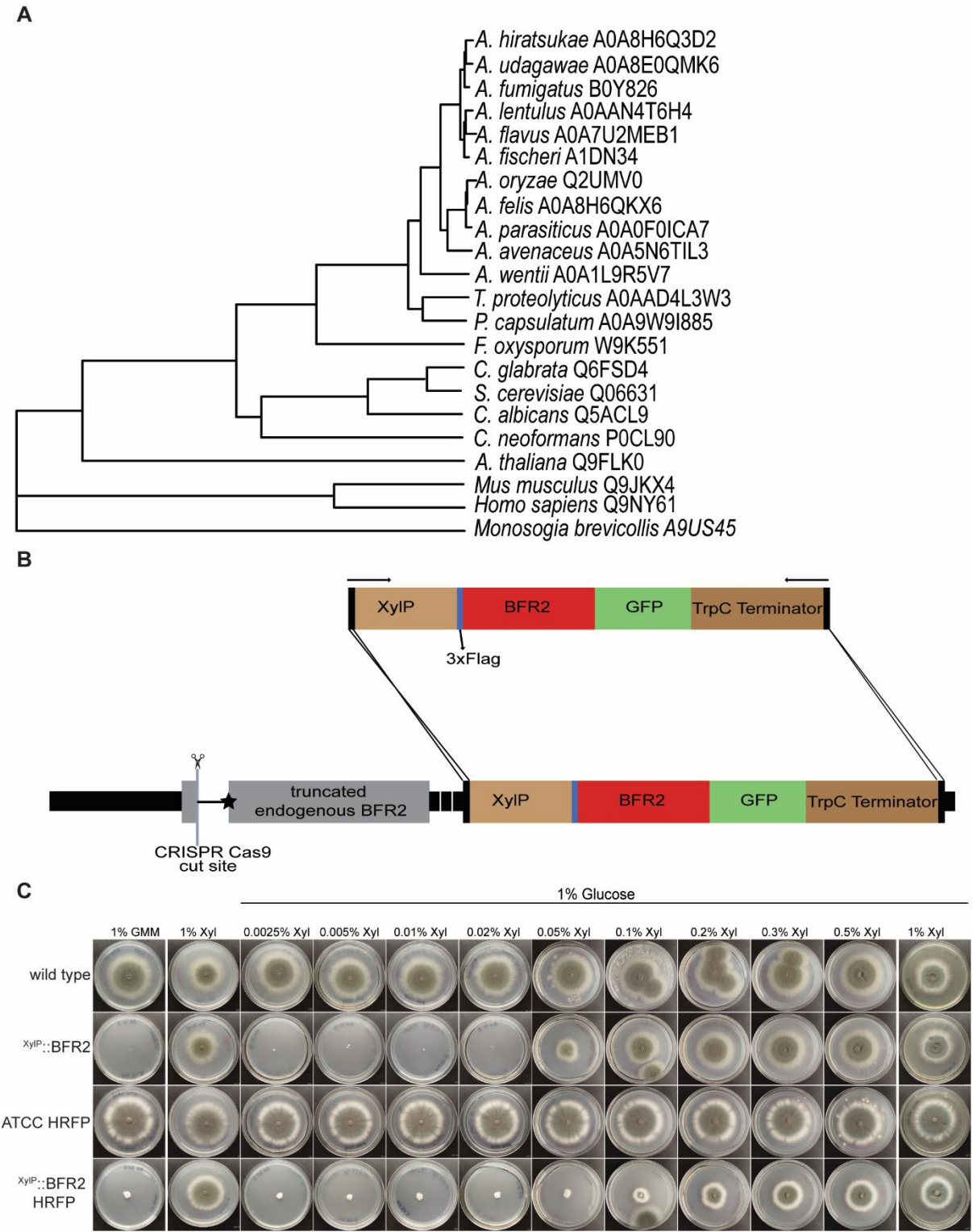

**Figure S1. BFR2 is a fungal homolog of mammalian AATF and is required for growth in *A. fumigatus*.** (A) Phylogenetic tree of BFR2/AATF homologs across fungi, plants, metazoa and a choanoflagellate outgroup. Sequences are identified by species and UniProt accession. Protein

sequences were aligned with [aligner, version] and the tree inferred by [method, substitution model, software] with [n] bootstrap replicates; the tree was rooted on *Monosiga brevicollis*. (B) Strategy for generating the conditional <sup>XylP::</sup>BFR2 strain. The endogenous BFR2 locus was cleaved by CRISPR-Cas9 (scissors) using a single guide RNA targeting a site near the translational start codon; repair by non-homologous end joining introduced an indel generating a premature stop codon (black star), leaving a truncated endogenous allele. A complementation cassette encoding 3xFlag-BFR2-GFP under the xylose-inducible xylP promoter, followed by the TrpC terminator, was introduced by microhomology-mediated CRISPR-Cas9 repair and integrated ectopically at an unmapped genomic locus. (C) Growth of the indicated *A. fumigatus* strains in wild type and HRFP backgrounds including <sup>XylP::</sup>BFR2 mutants strains in both backgrounds after 4 days at 37°C on minimal medium containing 1% glucose (GMM), 1% xylose, or 1% glucose supplemented with the indicated concentrations of xylose (0.0025–1%). Conidia were point-inoculated at the plate centre. Depletion of BFR2 on glucose abolishes growth, which is progressively restored with increasing xylose.

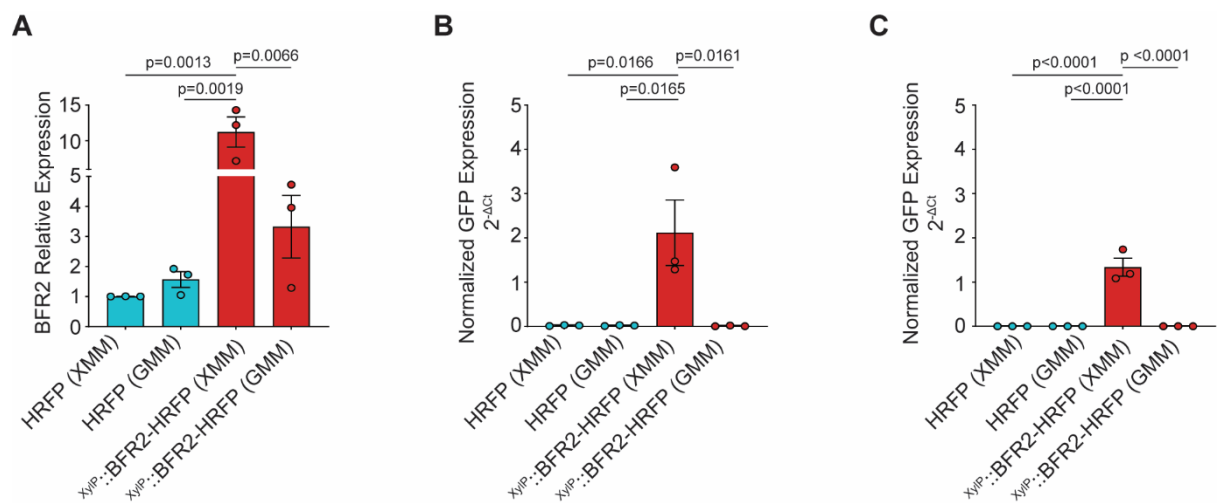

**Figure S2. Validation of conditional BFR2 expression in the HRFP background.** (A) BFR2 mRNA expression in HRFP and $XylP::BFR2$-HRFP strains following overnight growth of $5 \times 10^6$ conidia under fully inducing (1% XMM) or non-inducing (1% GMM) conditions at 37 °C. Expression was normalized to actin and is presented as mean relative BFR2 expression ($2^{-\Delta\Delta Ct}$) ± SEM from three independent biological experiments. Statistical significance was assessed by one-way ANOVA with Tukey's multiple comparisons test, with $p < 0.05$ considered statistically significant. (B–C) Relative expression of GFP (B) and FLAG (C), quantified as surrogate reporters of construct-specific transcript abundance under the same growth conditions. Because the GFP and FLAG sequences are present in the 3×FLAG-BFR2-GFP cassette in $XylP::BFR2$-HRFP but absent from the HRFP control strain, expression is reported as actin-normalized $2^{-\Delta Ct}$ values rather than $2^{-\Delta\Delta Ct}$ values. The HRFP control therefore provides no baseline Ct for a $\Delta\Delta Ct$ calculation. Data are presented as mean actin-normalized expression ± SEM from three independent biological experiments. Statistical significance was assessed by one-way ANOVA with Tukey's multiple comparisons test, with $p < 0.05$ considered statistically significant.
